# A simplified perchloric acid workflow with neutralization (PCA-N) for democratizing deep plasma proteomics at population scale

**DOI:** 10.1101/2025.03.24.645089

**Authors:** Vincent Albrecht, Johannes B. Müller-Reif, Vincenth Brennsteiner, Matthias Mann

**Affiliations:** Department for Proteomics and Signal Transduction, Max-Planck Institute of Biochemistry, Am Klopferspitz 18, 82152 Martinsried

**Author notes:** These authors contributed equally (Vincent Albrecht, Johannes B. Müller-Reif).

## Abstract

Large scale plasma proteomics studies offer tremendous potential for biomarker discovery but face significant challenges in balancing analytical depth, throughput and cost-effectiveness. We present an optimized perchloric acid-based workflow with neutralization – PCA-N – that addresses these limitations. By introducing a neutralization step following protein precipitation, PCA-N enables direct enzymatic digestion without additional purification steps, reducing sample volume requirements to only 5 µL of plasma while maintaining deep plasma proteome coverage. The streamlined protocol allows preparation of over 10,000 samples per day using 384-well formats at costs comparable to undepleted plasma analysis (NEAT). Rigorous validation according to the recently introduced CLSI C64 guideline demonstrated that despite somewhat higher technical variability compared to NEAT, PCA-N maintained excellent biological resolution and reproducibility. We confirmed the workflow’s exceptional stability through analysis of 1,500 quality control samples systematically interspersed among 36,000 plasma samples measured continuously over 311 days. Technical performance remained consistent across multiple instruments, sample preparation batches and nearly a year of measurements. Compared to NEAT plasma proteomics, PCA-N doubled the proteomic depth while maintaining comparable reagent costs and throughput. The minimal sample reequipments, operational simplicity while using only common laboratory chemical and exceptional scalability positions PCA-N as an attractive approach for population-level plasma proteomics, democratizing access to deep plasma proteome analysis.

**Highlights:**

- Perchloric acid with neutralization (PCA-N) allows drastic upscaling of plasma proteomics
- Minimal sample input (5 µL) in 384-well format enables thousands of samples processed daily
- PCA-N democratizes deep plasma proteomics using only common chemicals
- Validation per CLSI C64 demonstrates excellent repeatability and reproducibility
- PCA-N showed exceptional stability across 311 days of continuous MS acquisition

**Graphical Abstract:** 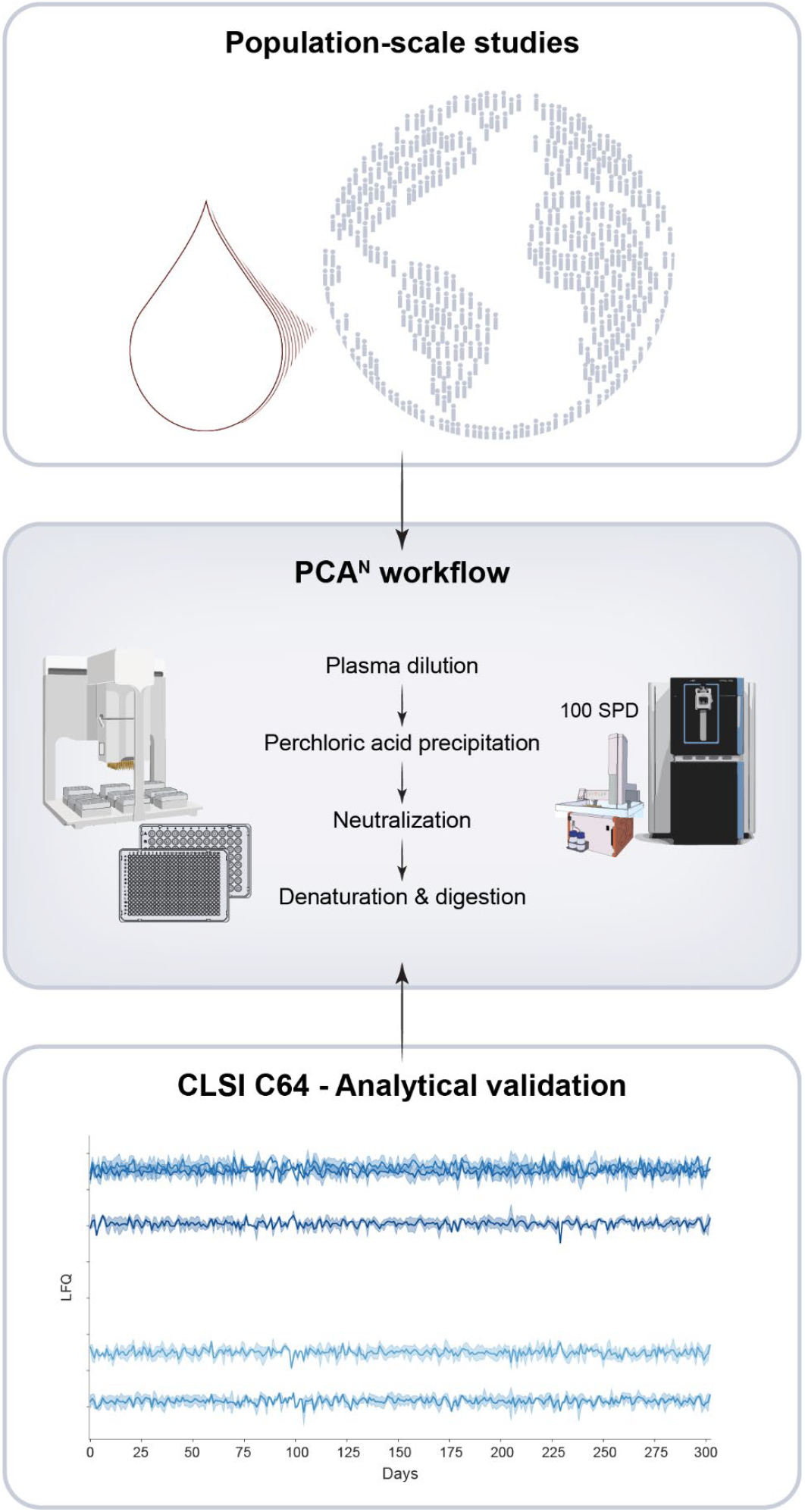

## Introduction

Plasma proteomics has undergone a remarkable transformation in recent years, evolving from small-scale exploratory studies to population-level analyses with tens of thousands of samples ^1–5^. This shift has been driven by the recognition that large cohorts are essential for robust biomarker discovery – enabling statistical power to identify subtle protein signatures while accounting for biological variability across populations^6^. Advancements in mass spectrometry (MS)-based plasma proteomics are a key enabler of this transformation^7–9^.

However, the field faces a critical tradeoff between analytical depth, robustness and practical feasibility for large cohorts. Standard undepleted (NEAT) plasma analysis offers simplicity and high throughput, but insufficient proteomic depth to detect low abundant, clinically relevant proteins. More elaborate techniques, such as antibody-based depletion^10,11^, nanoparticle enrichment^12–14^ and protein precipitation approaches^15,16^, may introduce practical limitations – including large sample volume requirements, high costs per sample, limited throughout, and challenges in long-term repeatability and susceptibility to varying sample quality – which become increasingly challenging as study sizes grow. These limitations also exist with affinity binder technologies, which despite successful application to population-scale studies through substantial investments^3,4^, remain cost-prohibitive in many research settings, creating a barrier to widespread plasma proteomic research at true population scale.

As studies continue to expand in size and scope, there is a need for workflows that can simultaneously deliver comprehensive, reliable proteome coverage and the operational efficiency required for extremely large cohorts^17^. With this challenge in mind, we here present a novel implementation of a perchloric acid (PCA)-based workflow – PCA-N, that overcomes these longstanding limitations. By incorporating a simple yet effective neutralization step, PCA-N eliminates the need for protein extraction, dramatically reducing sample volume requirements to just 5 µL of plasma and enabling parallelization to 384-well plates. This streamlined protocol allows preparation to an unparalleled scale of more than 10,000 samples per day at costs comparable to standard NEAT plasma analysis. This robust workflow overcomes long-standing barriers to extreme-scale plasma proteomics studies, making population-level studies technically and economically feasible.

We developed PCA-N specifically to enable preparation and analysis of extremely large cohorts and applied this methodology to an ongoing project involving 50,000 plasma samples. Here we describe PCA-N in detail and present an analytical validation for repeatability and reproducibility according to CLSI C64 guideline^18^. We also evaluate 1,500 quality control samples that we systematically included throughout a measurement campaign of approximately 36,000 plasma samples measured continuously over almost one year.

## Results

### Neutralization-based perchloric acid enrichment plasma proteomics workflow

The analysis of large cohorts calls for deep plasma proteome but also low marginal costs, high throughput and given the precious nature of clinical samples, minimal input volumes. Current deep plasma workflows relying on e.g. antibody-based depletion or bead-based enrichment have limitations in these critical aspects, hampering the scalability needed for ever increasing size of proteomic studies and widespread adoption of deep plasma workflows.

Perchloric acid (PCA) precipitation, originally developed many decades ago for glycoprotein analysis^19^, has recently seen new interest^20–22^. This technique exploits the differential solubility of plasma proteins in acidic conditions – high-abundant proteins including albumin and immunoglobulins precipitate, while many lower-abundance proteins remain in solution. The Steen laboratory recently harnessed this approach for modern plasma proteomics applications, developing a robust “perCA” workflow ^16^, and demonstrating its utility in large cohort analyses of over 3,000 samples with sufficient sensitivity to detect even viral proteins^23,15^.

Building upon this foundation, we found that one of the most critical steps is plasma protein digestion. The perCA protocol describes solid phase extraction (SPE) to desalt and re-buffer proteins from the PCA-containing supernatant before enzymatic digestion. Here, we introduce a critical innovation: a neutralization step that enables direct digestion of proteins in the supernatant without additional purification (**Figure 1A**). After PCA-mediated precipitation and centrifugation-based separation of the insoluble protein fraction, the soluble fraction is neutralized with sodium hydroxide (NaOH), yielding sodium perchlorate (NaClO_4_), the most water soluble of the common perchlorate salt^24^ (Eq 1).

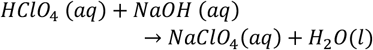

**Figure 1.**
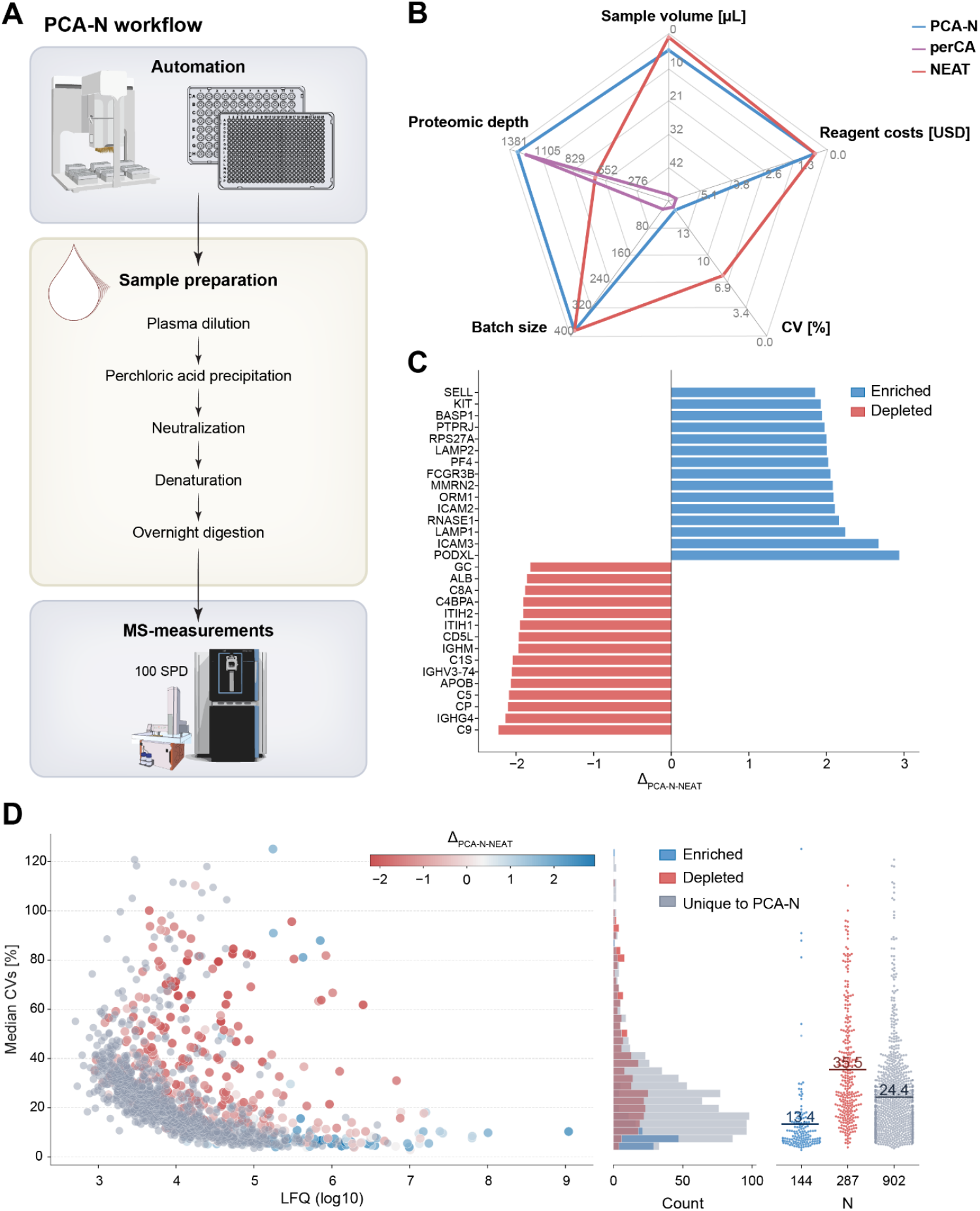
Neutralization-based perchloric acid enrichment plasma proteomics workflow for high throughput deep plasma proteomics. A. Schematic representation of the PCA N workflow. Plasma samples undergo PCA precipitation in 96 well format, followed by recovery of the supernatant containing enriched proteins. The neutralization step enables direct transfer to 384 well plates for denaturation and overnight digestion without clean-up, followed by high-throughput and high-sensitivity MS analysis at 100 samples per day (SPD). B. Radar plot comparing key performance metrics of PCA N, perCA and NEAT workflows, including sample volume requirements [µL], proteomic depth [protein groups identified], reagent costs [USD], coefficient of variation (CV) [%], and batch size. C. Waterfall plot showing the top 15 enriched (blue) and depleted (red) proteins by PCA N compared to NEAT, calculated as delta of intensities (Δ = PCA - NEAT). D. Relationship between protein abundance (log10 LFQ intensity) and measurement precision (CV [%]). Proteins are color coded as enriched by PCA N (blue), depleted by PCA N (red), or unique to by PCA N (gray). Right panel shows the distribution of enriched, depleted and unique proteins with their respective mean CVs [%].

This neutralization step reduces sample requirements compared to the published workflow by ten-fold – from 50 µL to just 5 µL of plasma (**Figure 1B**). In PCA-N 5 µL of plasma yield approximately 1000 ng of peptides after enrichment. With 200 ng desired per injection into the LC-MS system, the workflow could be further downscaled to use as little as 1 µL of plasma per analysis. The significant reduction in sample volume was made possible through the neutralization step, which allows loss-less recovery of proteins from the supernatant. This peptide yield is twice that of the SPE-based workflow, requiring only 1 µL of plasma for an equivalent MS injection compared to 2 µL for the previous method. The reduced volume requirements make the PCA-N workflow compatible with 384-well plates and potentially scalable to 1536-well formats. The original workflow is limited by the precipitation step and subsequent centrifugation, restricting throughput to 24 samples per batch in a standard bench-top centrifuge. The PCA-N workflow thus increases throughput up to 16-fold. Through strategic workflow scheduling, PCA-N can facilitate preparation of up to 13,000 samples per day in a highly parallelized fashion (**Figure S 1**.).

The preparation cost per sample decreases correspondingly, from about 5 USD for cleanup – mainly due to the cost of the plates – to the negligible cost of the traditional NEAT plasma preparation, as only digestion reagents need to be accounted for. In both workflows, the main remaining cost factor isthe digestion enzymes, making the overall sample processing expenses comparable. We found that omitting the µSPE-HLB plates has minimal impact on analytical performance, with similar mean coefficient of variation (CVs) (16.1 % for PCA-N versus 16.4 % for perCA) and comparable proteomic depth (1315 proteins in PCA-N versus 1239 proteins in perCA and 2509/2529 overall for five technical samples, see below) (**Figure 1B**). Note that proteomic depth is highly dependent on the plasma and in this benchmarking study, we used high-quality highly purified plasma samples, devoid of platelet and other contamination. Using these same samples, the NEAT workflow identified 639 protein groups per sample (781 proteins overall) using optimized DIA settings (Experimental Methods, **Figure S 2**.). Based on our experience with various clinical cohorts (see below) and their inherent differences in sample quality, the PCA-N workflow can easily identify 2,000 proteins per sample, with cumulative protein identifications exceeding 4,000 for larger cohorts (> 1,000 samples).

The PCA-N workflow thus combines the advantages of NEAT plasma profiling (reagent costs for preparation of under 1 USD per sample, high throughput capacity with 384-well plate batches), while doubling the proteomic depth (**Figure 1B**). We conclude that PCA-N makes robust deep plasma proteomics accessible for large-scale clinical applications.

### Proteins enriched and depleted by perchloric acid treatment

The increased depth of the PCA plasma workflow is attributed to a more balanced dynamic range, likely resulting from the preferential depletion of the most abundant proteins. Major depleted proteins include albumin, complement component 9 (C9), ceruloplasmin (CP) and inter-alpha-trypsin inhibitor heavy chain H2 (ITIH2)^16^,which our data confirms (**Figure 1C**). To systematically characterize the specificity of PCA-based protein enrichment or depletion, we calculated a delta in protein intensity for proteins identified in the NEAT and PCA-N workflow (Δ = PCA-N - NEAT). Proteins with Δ>0 were classified as enriched and those with Δ<0 as depleted (**Figure 1C**). While only 431 proteins were detected in both workflows, a striking pattern emerged with respect to the precision of quantification. Enriched proteins exhibited substantially improved technical reproducibility, with a mean CV of 13.4 % compared 36.5 % for depleted proteins. This observation explains why the PCA-N workflow (like most enrichment techniques) shows higher overall technical variability while still providing robust biological information – the variability is primarily associated with depleted high-abundance proteins rather than the enriched lower-abundance proteins that drive biological insight. This differential effect on proteins is further illustrated by the relationship between measurement precision and protein abundance. Notably, a substantially larger fraction of proteins was uniquely detected in the PCA-N workflow rather than uniquely identified in NEAT (67.7 % vs 34.9 %), highlighting its superior sensitivity for detecting less abundant plasma proteins (**Figure 1D**).

### Benchmarking the PCA-N workflow based on the CLSI C64 guideline

To rigorously validate the PCA-N workflow for potential clinical applications, we conducted a comprehensive evaluation following the Clinical and Laboratory Standards Institute (CLSI) C64 guideline for assessing reproducibility and repeatability^18^. First published in 2021 and termed *Quantitative Measurement of Proteins and Peptides by Mass Spectrometry*, this standardized framework provides a systematic framework for developing clinical protein and peptide assays from conception to validation, particularly for those who have experience with traditional small-molecule liquid chromatography-mass spectrometry (LC-MS) but not with protein and peptide analysis. Recent literature strongly emphasizes the importance of following international analytical guidelines when developing MS-based assays for peptide quantification^25–27^. To our knowledge, our study represents the first application of CLSI C64 to evaluate the reproducibility and repeatability of discovery plasma proteomics workflows.

As a reference to the PCA-N workflow we conducted a head-to-head comparison with the NEAT workflow. We prepared five biological (plasma from different individuals) and five technical replicates (identical aliquots of pooled quality control plasma) five times per day over five consecutive days, followed by sequential MS measurements. Additionally, the protocol was extended to assess repeatability over 20 consecutive days with two replicates per specimen (**Figure 2A**).

**Figure 2.**
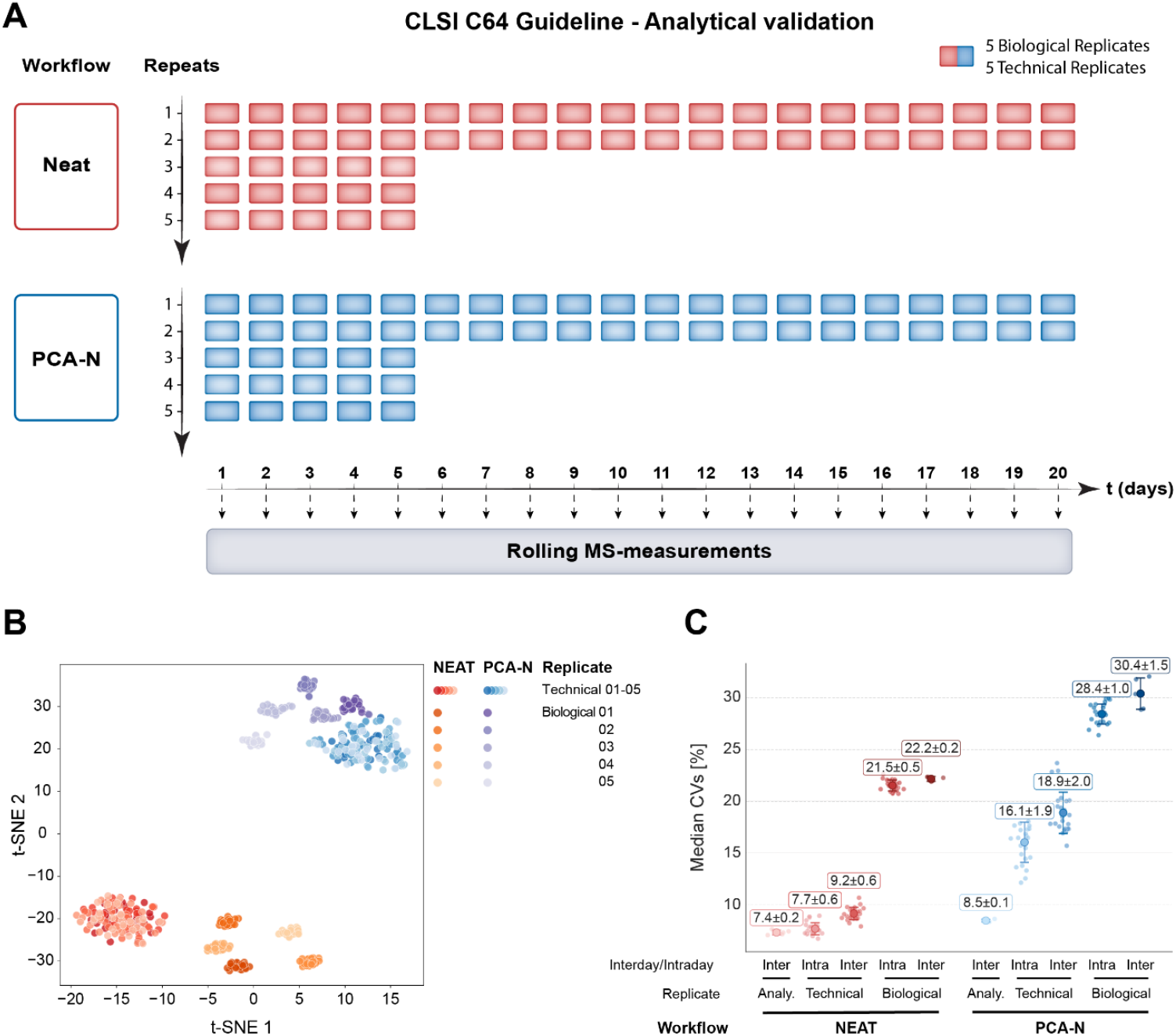
Comprehensive validation of the PCA-N workflow according to CLSI C64 guidelines. A. Study design for assessing reproducibility and repeatability according to CLSI C64 guidelines. Five biological replicates and five technical replicates were prepared with both NEAT (red) and PCA N (blue) workflows with five repeats per sample for the first five days, followed by two repeats per sample for days 6 – 20. MS measurements were performed in rolling fashion after one month of peptide storage. B. t-SNE visualization of all samples showing clear separation between NEAT (red/orange shades) and PCA N (blue/purple shades) workflows. Technical and biological replicates form distinct clusters, demonstrating both technical reproducibility and biological resolution. C. Multi-level analysis of coefficient of variation (CV) to assess variance at different stages: analytical (re-loading of peptides on Evotips and LC-MS performance), technical (sample preparation replicates) and biological (from five individuals). Variability is evaluated under both intraday and interday conditions for both workflows. CV values are calculated based on median calculations and presented as mean CV [%], with standard deviations indicated by error bars.

### Reproducibility

Dimensionality reduction using t-SNE (**Figure 2B**) clearly separated the PCA-N and NEAT results, and distinctly clustered technical and biological replicates within each workflow. Reproducibility was systematically evaluated using a multi-tiered CV calculation approach (**Figure 2C**). Analytical reproducibility – concerning peptide loading, LC-separation and MS-performance – was assessed by repeatedly measuring pooled peptides for each workflow. Additionally, we used technical and biological samples to calculate both intraday and interday CVs to evaluate workflow reproducibility and biological/clinical resolution. We observed a systematic increase in technical variation over time for both workflows. The NEAT workflow had with CVs of 7.7|9.2 % (intraday|interday) excellent technical reproducibility, comparable to its analytical variability of 7.4 %. The PCA-N workflow showed approximately twice the technical variability, with CVs of 16.1|18.9 %. Importantly, biological CVs scaled proportionally, with 21.5|22.2 % for NEAT and 28.4|30.4 % for PCA-N (intraday|interday). These values represent an additional biological variation component of 13.8|13 % for NEAT and 12.3|11.5 % for PCA-N.

### Repeatability

The extended timeframe of 20 consecutive days of sample preparation and MS measurements enabled a comprehensive evaluation of long-term stability under conditions mimicking routine clinical laboratory operations. To evaluate this, we categorized quantified proteins into quartiles by their CV values and assessed consistency of ranking in the reproducibility experiment and 20-day interday measurements (**Figure 3A, 3D**). This analysis revealed that 68 % of proteins in the NEAT workflow and 65 % of proteins in the PCA-N workflow maintained consistent CV rank categories (**Figure 3B, 3E**). This remarkable similarity in stability profiles demonstrates that despite higher absolute CV values, the PCA-N workflow provides repeatability comparable with the conventional NEAT approach, but with twice the proteomic depth. This finding is also confirmed by Pearson correlations of 0.90 for NEAT and 0.88 for PCA (of log_10_-transformed intraplate and interplate CVs) (**Figure S 3A, S 3B**), indicating highly consistent relative quantification over the extended timeframe. Most importantly, when assessing biological samples over the 20-day period, sample-to-sample correlation analysis demonstrated robust separation of individual biological samples (**Figure 3C, 3F**). Collectively, these results demonstrate that the PCA-N workflow, while showing higher technical variability compared to NEAT, maintains excellent repeatability characteristics.

**Figure 3.**
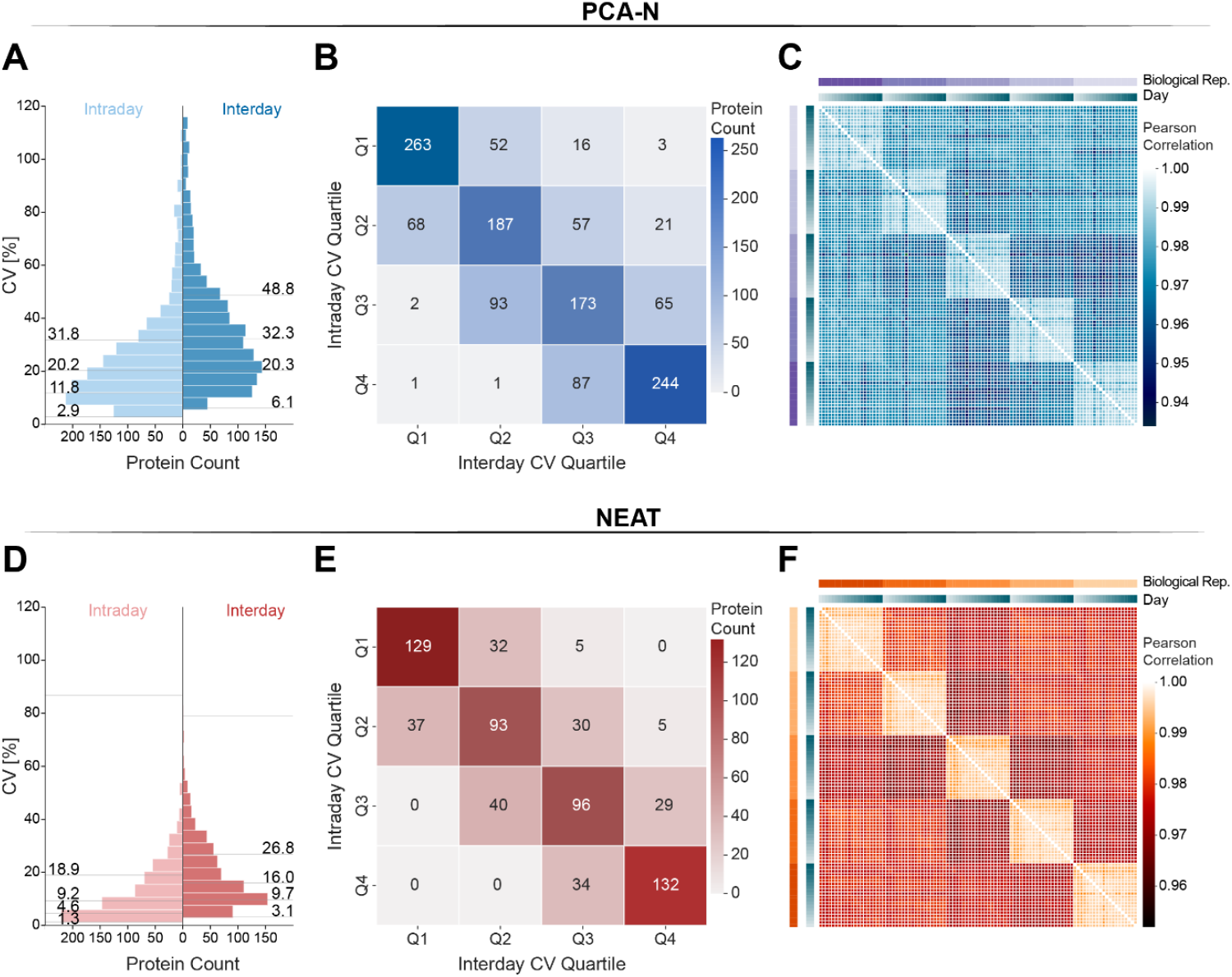
Long term repeatability assessment of PCA-N (A, B, C) and NEAT (D, E, F) workflows according to the CLSI C64 guideline. A, D. Distribution of median CV values for proteins in both intraday (light blue/red, derived from reproducibility experiment) and interday (dark blue/red) measurements. Numbers indicate median CVs [%] for each distribution quartile (Neat: N = 662, Stability = 68 %, PCA: N = 1,333, Stability = 65 %). B, E. Confusion matrices showing the stability of protein CV rankings between intraday and interday measurements. Numbers in each cell represent protein counts, with diagonal cells indicating proteins maintaining the same quartile rankling. C, F. Sample to sample correlation heatmaps for biological replicates over 20 days of measurements. Color intensity represents correlation coefficient values and distinct clustering patterns are visible for samples from the five individuals. Biological replicate and day information is indicated on the top and right of each heatmap.

#### Long-term stability assessment in an extremely large-scale proteomics study

We specifically developed the PCA-N workflow to enable preparation and analysis of extremely large cohorts. This methodology is currently applied to an ongoing large-scale project involving 50,000 plasma samples of the Multi-Omics for Mothers and Infants (MOMI) Consortium ^17,28^. While the clinical findings of that multi-center study are not the subject of this report, we present a comprehensive technical evaluation of the 1,500-quality control (QC) plasma samples that we prepared and measured as every 24^th^ sample, representing a total of 36,000 LC-MS runs and providing a unique opportunity to assess long-term stability under real-world conditions. Total preparation time for the individual MOMI cohorts was just six days (sample preparation batches) in total. For each cohort we prepared the QC samples using the optimized and parallelized PCA-N workflow, with measurements conducted for almost one year on two Orbitrap Astral mass spectrometers working in parallel.

Non-linear dimensionality reduction using t-SNE revealed that mass spectrometers drove primary separation between samples (MS #1 versus MS #2, **Figure 4.A**), while no separation was observed between different batches within each instrument cluster, confirming excellent reproducibility across sample preparation batches. After ComBat-based batch correction^29^ for the mass spectrometers, neither the instrumentation nor the batches caused separation (**Figure 4.B**). The intraplate CVs assessing the technical variability across all QC samples was 16.0 % (overall mean of the median CVs) (**Figure 4C**., **Figure S 4A**). This value is remarkably consistent with the intraday CVs observed in our CLSI C64 reproducibility benchmarking experiments (**Figure 2.C**). Individual cohort batches showed a similar pattern, with intraplate CVs ranging from 15.8 % to 16.9 %, further demonstrating robust reproducibly of the PCA-N workflow across different sample batches and extended measurement periods. Long-term stability of protein quantification is exemplified by the top five proteins with the lowest CVs which demonstrated very high stability throughout the nearly year-long measurement period (**Figure 4.G**, **Figure S 4E**). Long-term repeatability was then evaluated using the quantile-based stability analysis, where QC samples were randomly interspersed throughout the measurement of biological samples. This revealed that 65 % of proteins maintained consistent CV rank categories between intraplate and interplate measurements (**Figure S 4C**), identical to the intraday stability score observed in our controlled CLSI C64 experiments. The interplate Pearson correlation was 0.890 (**Figure S 4D**), indicating excellent quantitative consistency across the entire measurement period of almost one year. After batch correction, despite slightly higher intraplate CVs of 17.7 % (**Figure 4.C**), stability further increased to 81.5 % and Pearson correlation to 0.97 (**Figure 4.D-4.F**).

**Figure 4.**
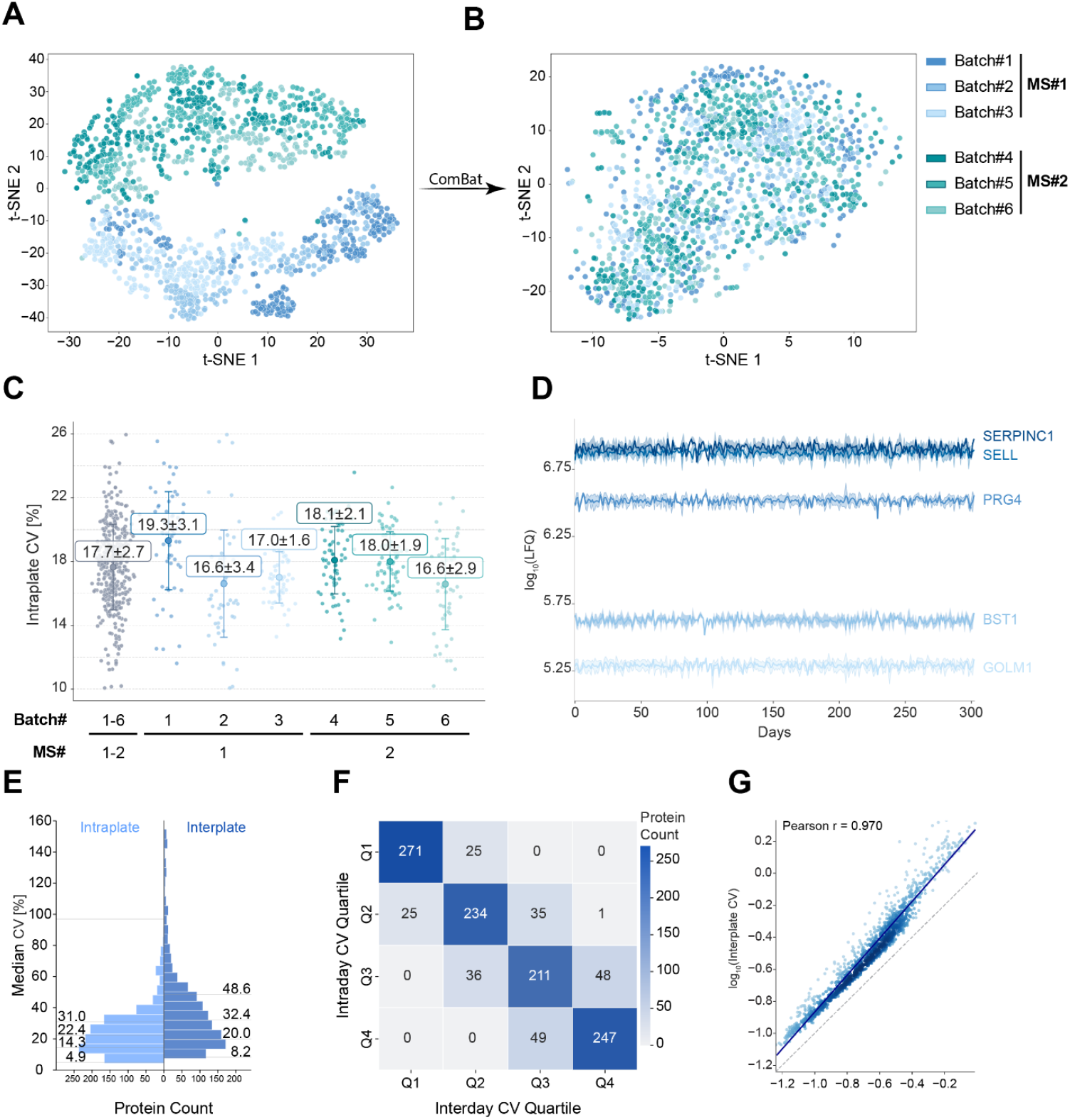
Technical validation of the PCA-N workflow in an extreme scale plasma proteomics study. A, B. t SNE visualization of 1,500 quality control plasma samples from six sample preparation batches measured on two mass spectrometers (MS #1 and MS #2) over 311 days as every 24th sample as part of a 50,000-sample study. Without batch correction samples cluster primarily by MS instrument rather than cohort, demonstrating consistent sample preparation across batches. After batch correction, neither MS instrument nor sample batches drive separation. Batch corrected data is used for the following displays. C. Intraplate median coefficients of variation (CVs) for the combined dataset and split by the individual sample batches/cohorts. D. Reproducibility of the label-free quantification (LFQ) intensities of the top five proteins with the lowest CVs throughout the 311-day measurement period. The line represents the mean values and the shading the standard deviation for SERPINC1 (Antithrombin-III) (CV_SERPINC1_ = 7.0 %), SELL (L-selectin) (CV_SELL_ = 6.1 %), PRG4 (Proteoglycan 4) (CV_PRG4_ = 5.6 %), BST1 (ADP-ribosyl cyclase/cyclic ADP-ribose hydrolase 2) (CV_BST1_ = 7.2 %), GOLM1 (Golgi membrane protein 1) (CV_GOLM1_=7.1 %). E. Distribution of CV values for proteins in both intraplate (light blue) and interplate (dark blue) measurements (N=1,500). Numbers indicate median CVs [%] for each distribution quartile. F. Confusion matrices showing the stability of protein CV rankings between intraplate and interplate measurements. Numbers in each cell represent protein counts, with 81.5 % of proteins maintaining the same quartile rankling (diagonal values). G. Pearson correlation of log10-transformed intraplate and interplate CVs.

These results validate the robustness and scalability of the PCA-N workflow for extreme high-throughput applications. The ability to maintain consistent technical performance across thousands of samples, multiple instruments and an extended measurement period validates PCA-N as a reliable platform for large-scale plasma protomeric studies.

## Discussion

The analysis of large clinical cohorts requires plasma proteomics workflows that deliver deep proteome coverage while maintaining cost-effectiveness, high-throughout and minimal sample consumption. The PCA-N workflow presented here accomplishes these objectives by a neutralization step that builds upon and significantly enhances the previously developed perchloric acid precipitation workflow. By eliminating the need for solid phase extraction and enabling direct enzymatic digestion after neutralization, this workflow substantially reduces sample requirements from 50 µL to just 5 µL of plasma, with the potential for further downscaling to as little as 1 µL. The minimal sample volume requirement facilitates automation and compatibility with 384-well and potentially 1536-well formats, substantially increasing throughput capacity compared to conventional workflows.

When benchmarked against the standard NEAT plasma workflow, PCA-N doubles the proteome coverage, while maintaining similar reagent costs under 1 USD per sample. This combination of depth, throughout, and cost-effectiveness positions PCA-N as an attractive approach for extreme-scale plasma proteomics studies, even in resource-limited settings.

Our rigorous validation according to CLSI C64 guideline revealed that despite somewhat higher technical variability in PCA-N compared to NEAT plasma analysis, both workflows show comparable resolution between clinical samples. This biological component of the total CV was remarkable similar, indicating that disease-relevant proteomic changes can be captured effectively despite the increased technical variability inherent to the depletion process. Long-term repeatability assessment demonstrated that both workflows maintain similar stability profiles over extended periods. Despite higher absolute CVs, PCA-N provides comparable repeatability to the conventional NEAT approach while delivering twice the proteome depth. This consistency was further confirmed through robust separation of biological samples over a 20-day period, which is particularly crucial for large-scale clinical studies, where sample acquisition an analysis may occur over weeks or months. In conclusion, the remarkable consistency of intraplate CVs across different cohort batches and high interplate correlation demonstrate exceptional technical performance across thousands of samples and extended measurement periods. The successful application of PCA-N to over 34,500 plasma sample across 311 days validates its suitability for large-scale clinical studies. This establishes a new paradigm for large-scale studies that advances biomarker discovery and validation by MS-based proteomics. By drastically reducing sample volumes and costs while maintaining excellent proteome depth and reproducibility, this approach enables population-scale studies previously considered unfeasible due to technical or resource constraints. Despite these advances, we recognize opportunities for continued refinement. These include optimized enzymatic digestion protocols, internal standard spike-in procedures for absolute quantification and alternative depletion/enrichment strategies that could further enhance the capabilities presented here. Together, such developments are transforming our ability to identify novel biomarkers and disease mechanisms across large, diverse populations.

## Acknowledgement

We thank members of the Steen2 Lab for generously sharing their expertise and providing support in implementing the perCA workflow. We also acknowledge support by our colleagues, especially Igor Paron and Tim Heymann for their assistance with mass spectrometry measurements and data acquisition. The feedback and discussions with members of the Mann Labs, including Philipp Geyer, significantly improved this work. This work is supported by the Max Planck Society for Advancement of Science and by the Bill & Melinda Gates Foundation (INV-058348).

## Supplementary Material

**Figure S1.**
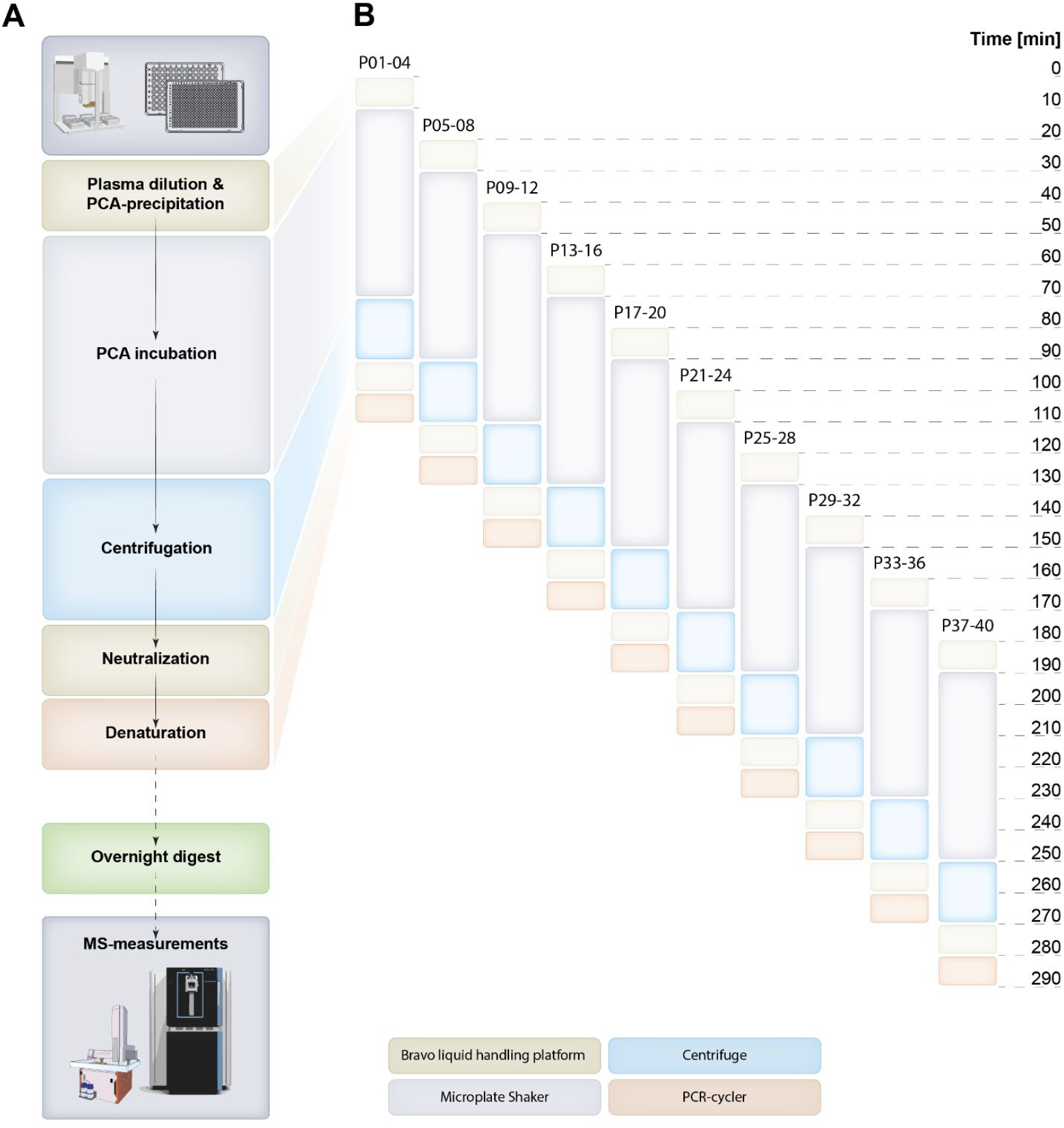
PCA-N workflow enables highly parallelized sample preparation. **A**. Schematic representation of the PCA-N workflow with main processing steps shown in sequential order from plasma sample to mass spectrometry analysis. **B**. Timeline illustrating the parallelized sample prepration strategy. The workflow leverages staggered use of liquid handling systems, centrifuges, and PCR-cycler to efficiently process multiple plates simultaneously. This example demonstrates the processing of 40 96-well plasma plates (in batches of 4 plates) in under 300 min.

**Figure S2.**
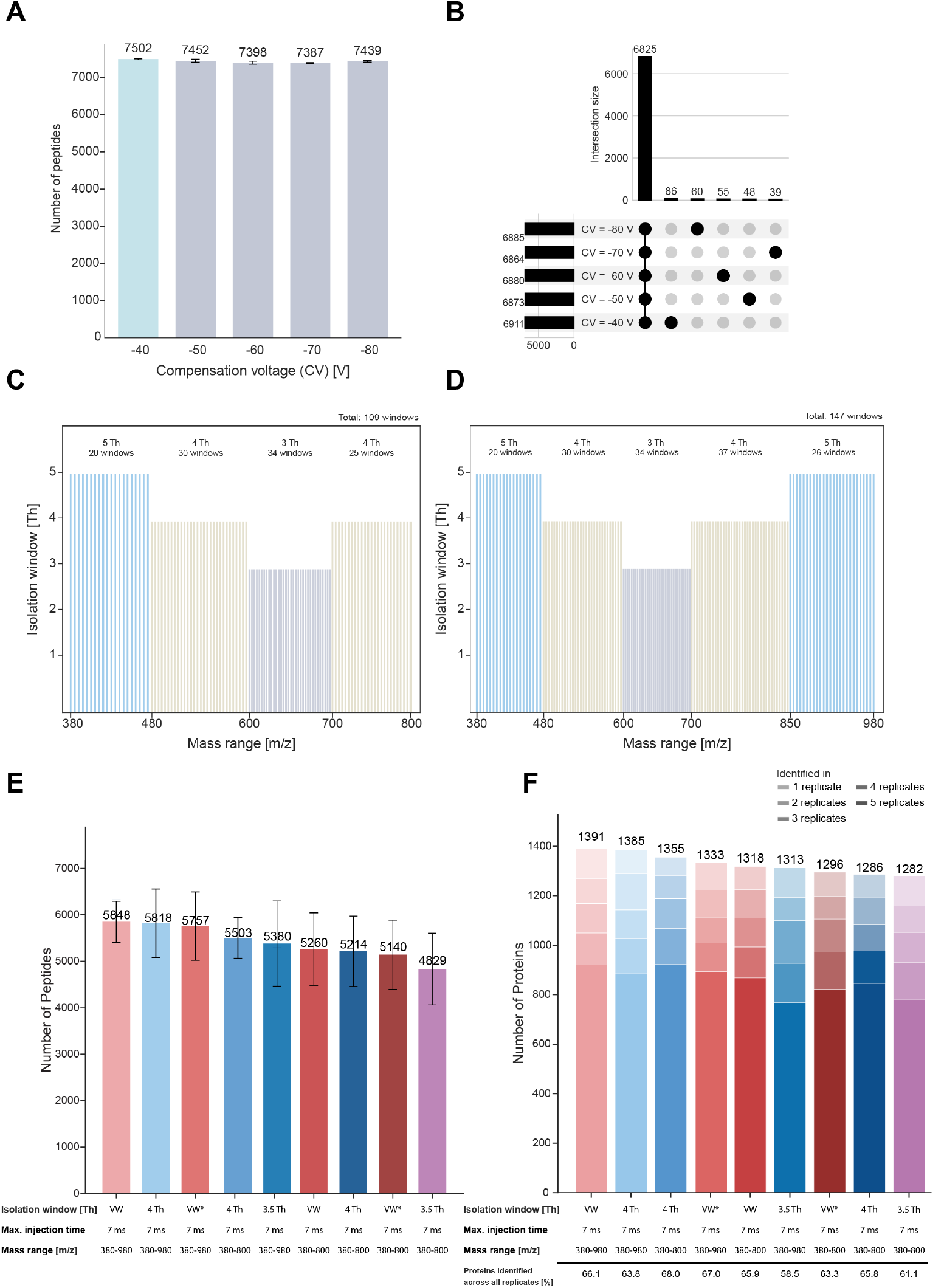
Mass spectrometry method optimization for deep plasma proteome coverage. A. Number of peptide identifications using different FAIMS compensation voltages (CV) ranging from −40 to −80 V. B. UpSet plot showing unique peptides per CV and the intersection of peptides shared across multiple CV settings. Other overlaps are not shown. C. Variable isolation window method with 5, 4, 3 and 4 Th windows spanning mass range 380-800 m/z (total 109 windows). D. Extended mass range variable isolation window method with 5, 4, 3, 4 and 5 Th windows spanning mass range 380-980 m/z (total 147 windows). E. Number of peptides identified using different MS methods comparing standard equally spaced windows and variable width windows (VW) (VW* = using optimal window placement). F. Number of proteins identified across 1 - 5 analytical replicates using different MS methods, with colour intensity indicating the number of replicates in which proteins were detected. Numbers above bars indicate total proteins identified. Percentage value show identification rates across all replicates.

**Figure S3.**
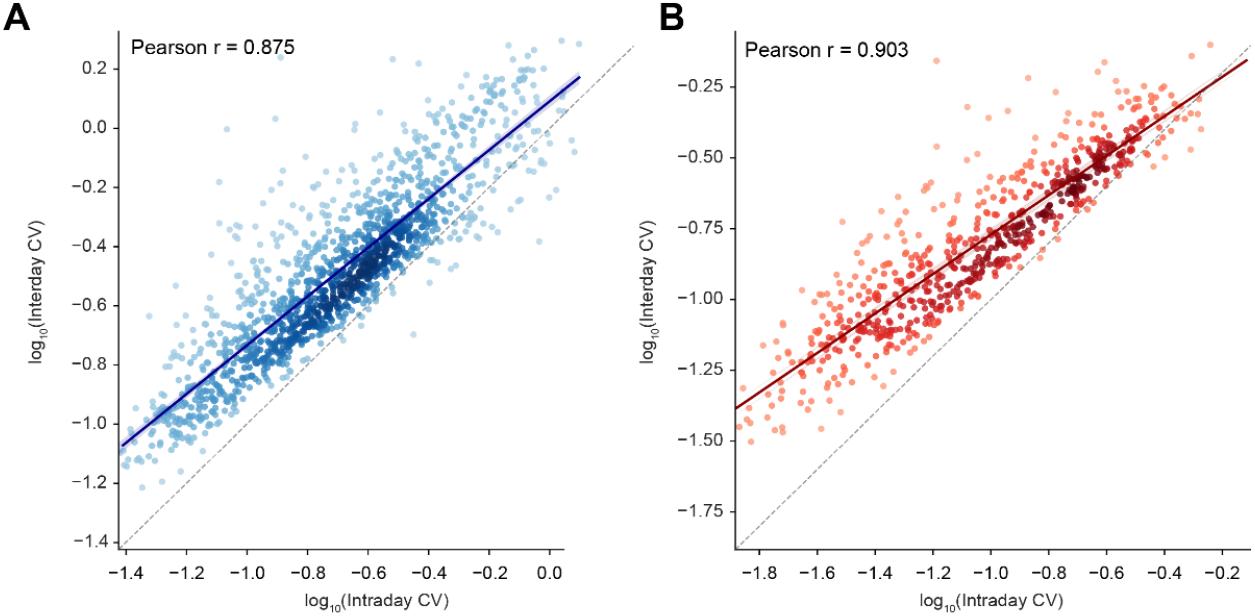
Long term repeatability assessment of PCA-N and NEAT workflows according to the CLSI C64 guideline. Pearson correlation of log10-transformed intraplate and interplate CVs for PCA N (A) and NEAT (B).

**Figure S4.**
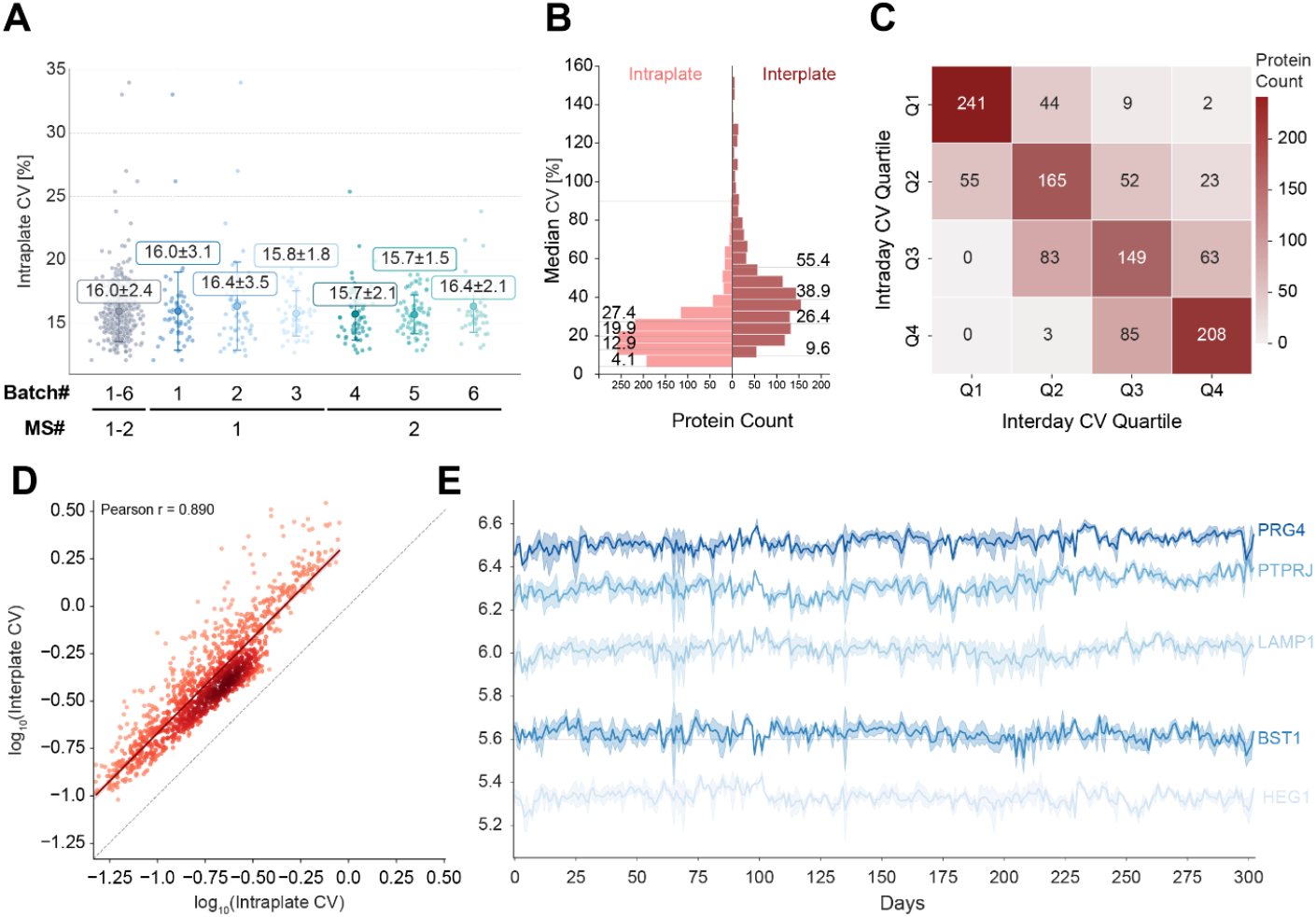
Technical validation of the PCA-N workflow in an extreme scale plasma proteomics study. Non-batch corrected data is used for the following displays. A. Intraplate median coefficients of variation (CVs) for the combined dataset and split by the individual sample batches/cohorts. B. Distribution of CV values for proteins in both intraplate (light red) and interplate (dark red) measurements (N=1,500). Numbers indicate median CVs [%] for each distribution quartile. C. Confusion matrices showing the stability of protein CV rankings between intraplate and interplate measurements. Numbers in each cell represent protein counts, with 65 % of proteins maintaining the same quartile rankling (diagonal values). D. Pearson correlation of log10-transformed intraplate and interplate CVs. E. Reproducibility of the label-free quantification (LFQ) intensities of the top five proteins with the lowest CVs throughout the 311 day measurement period. The line represents the mean values and the shading the standard deviation for PRG4 (Proteoglycan 4) (CV_PRG4_ = 7.4 %), BST1 (ADP-ribosyl cyclase/cyclic ADP-ribose hydrolase 2) (CV_BST1_ = 8.7 %), PTPRJ (Receptor-type tyrosine-protein phosphatase eta) (CV_PTPRJ_ = 8.8 %), LAMP1 (Lysosome-associated membrane glycoprotein 1) (CV_LAMP1_ = 8.8 %) and HEG1 (Protein HEG homolog 1) (CV_HEG1_ = 8.9 %).

## Experimental procedures

### Plasma Collection from Healthy Donors

Plasma was obtained by collecting blood in EDTA tubes (BD Vacutainer K2E Ref 367525 [10 mL; 18 mg EDTA]). After collection, blood was gently mixed by inverting the tube three times and centrifuged at 2000 x g for 20 min at 4 °C. The plasma layer was carefully collected without disturbing the buffy coat layer, aliquoted, snap-frozen, and stored at −80 °C. Blood was sampled from five healthy donors who provided written informed consent, with prior approval of the ethics committee of the Max Planck Society for the Advancement of Science in accordance with the Declaration of Helsinki principles.

For CLSI C64 guideline implementation, biological plasma samples were prepared and analyzed separately, while for the technical replicates, plasma samples from 5 individuals were pooled.

### Sample Preparation

#### NEAT Workflow

The NEAT workflow refers to a standard plasma proteomic workflow without depletion or enrichment. All subsequent steps were semiautomated using the Bravo liquid handling platform (Agilent). Briefly, 1 µL plasma was mixed with tris(2-carboxyethyl)phosphine (TCEP, C_final_=10 mM) and 2-chloracetamid (CAA, C_final_ = 40 mM) in a Tris buffer (pH 8.5, C_final_ = 100 mM) in a 384-well plate. Proteins were reduced, alkylated and denatured at 95 °C for 10 min in a PCR cycler. Digestion was performed overnight at 37 °C using trypsin (0.5 µg/µL, 1 µL) and LysC (0.5 µg/µL, 1 µL). The digest was stopped with trifluoroacetic acid (TFA, C_final_ = 1 %).

#### PCA-N Workflow

The PCA-N workflow was semiautomated using the Bravo liquid handling platform (Agilent). Five µL plasma were diluted in water (25 µL),mixed with perchloric acid (PCA, 1 M, 25 µL, C_final_ = 0.5 M) and incubated at 4 °C for 60 min in 96-well plates. The suspension was centrifuged at 4000 x g for 20 min at 4 °C, following transfer of supernatant (24 µL) to a 384-well plate and adjustment to pH 8 - 8.5 using precisely titrated sodium hydroxide solution (NaOH, 1.4 M, 8 µL) to provide optimal conditions for enzymatic digestion. Subsequently, proteins were reduced, alkylated and denatured at 95 °C for 10 min using dithiothreitol (DTT, C_final_=10 mM) and CAA (C_final_ = 40 mM) in a triethylammonium bicarbonate buffer (TEAB, pH 8-8.5, C_final_ = 60 mM). n-Dodecyl β-D-maltosid (DDM, C_final_ = 0.01 %) was used as detergent ensuring compatibility with subsequent C18-based peptide desalting. Proteins were digested using trypsin/LysC (each 0.125 µg/µL, 0.8 µL) and digestion was stopped with TFA (C_final_ = 0.5 %).

#### perCA Workflow

The PCA-N workflow was benchmarked against the previously published perCA method^15,16^. In brief, fifty µL plasma were diluted with water (450 µL) (C_final_ = 2.5-3 µg/µL) in 1.5 mL tubes, mixed with concentrated PCA (11.7 M, 25 µL, C_final_ = 0.56 M) and incubated at −20 °C for 15 min. The suspension was centrifuged for 60 min at 4 °C at 3200 x g using an Eppendorf 5425 R. The supernatant (390 µL) was acidified with TFA (40 µL, 1 %, C_final_ = 0.1 %) and desalted on a μSPE-HLB plate (Oasis® HLB μElution Plate, Waters Corporation) according to manufacturer’s instructions to remove the PCA. Eluted proteins were digested as described for MS analysis.

### LC-MS Sample Preparation and Analysis

For all workflows, 200 ng of the digested peptides were loaded onto disposable Evotip C18 trap columns (Evosep Biosystems) according to the manufacturer’s instructions using the Bravo liquid handling platform (Agilent). In brief, Evotips were activated in 1-propanol, washed with 0.1% formic acid (FA) in acetonitrile (ACN) (1 min, 700 x g) and soaked again in 1-propanol. Tips were washed and then equilibrated with 0.1 % FA in water (Buffer A). After loading samples (3 min, 700 x g), tips were washed with Buffer A and maintained with 180 µL Buffer A to prevent drying. Tips were stored at 4 °C if not measured immediately.

Mass spectrometric analysis was performed using an Evosep One liquid chromatography system (Evosep Biosystems)^30^ coupled to an Orbitrap Astral mass spectrometer (Thermo Fisher Scientific) operated in Data Independent Acquisition (DIA) mode. Peptides were separated over 11.5 min on an 8 cm, 150 µm ID Aurora Rapid column (IonOpticks) using the Evosep 100 samples per day (SPD) method. The column was operated at 50 °C and interfaced with an EASY-Spray Source with a spray voltage of 1900 V.

The Astral mass spectrometer was interfaced with a FAIMS Pro device operated with a total carrier gas flow of 3.5 L/min and a compensation voltage of −40 V. MS1 scans (380 – 980 m/z) were acquired at a resolution of 120,000 with an AGC target of 500 % and a maximum injection time of 3 ms. MSMS scans were acquired in the Astral analyzer with 4 Th windows (scan range 150 - 2000 m/z), a maximum injection time of 7 ms and an AGC target of 500 %. HCD fragmentation was performed with 25% normalized collision energy.

### Data Analysis and Statistical Evaluation

Raw data was analyzed using DIA-NN 1.8.1 with an *in silico* predicted library on a high-performance computing cluster searching against the human TrEMBL FASTA database (taxonomy ID 9606) downloaded from UniProt in November 2023. Mass accuracy and MS1 accuracy were set to 10 ppm with a scan window radius of 6.

#### Quantitative assessment of technical variation

Reproducibility was systematically evaluated using a multi-layered coefficient of variation (CV) approach:

### Intraday analytical reproducibility

CVs were calculated across analytical replicates within the same day of sample preparation and acquisition to assess analytical variability introduced by sample loading, LC and MS performance.

### Intraday technical reproducibility

CVs were calculated across technical replicates within the same day of sample preparation and acquisition to assess sample preparation consistency within a batch.

### Interday technical reproducibility

CVs were calculated within technical replicates across days of sample preparation and acquisition to assess sample preparation consistencies across batches.

### Intraday biological reproducibility

CVs were calculated across biological samples within the same day of sample preparation and acquisition to quantify the individual-specific variation within a batch.

### Interday biological reproducibility

CVs were calculated within biological samples across days of sample preparation and acquisition to quantify the individual-specific variation across batches.

#### Depletion/Enrichment Analysis

Differences between the PCA-N and NEAT sample preparation were quantified by calculating the delta value (Δ_PCA-NEAT_) for each protein detected in both conditions. Proteins were classified as enriched (Δ>0) or depleted (Δ<0) in PCA-N compared to NEAT, enabling the identification of workflow-specific quantification biases.

#### Repeatability analysis

To evaluate whether proteins maintain consistent measurement precision across experimental conditions, a rank stability analysis was implemented. Proteins were categorized into quartiles based on their CV values, and the concordance between intraday and interday rankings was visualized using confusion matrices. We defined a stability percentage as the proportion of proteins remaining in the same rank category across both assessment levels. Pearson correlation coefficients were calculated to quantify the relationship between intraday/intraplate and interday/interplate CV measurements.

#### Dimensionality reduction

Prior to dimensionality reduction, missing values were imputed using K-nearest neighbors (KNN) algorithm (sklearn). A minimum valid value threshold of 5% across samples was applied to filter low-confidence proteins. To capture complex, non-linear relationships in the proteomic data, t-Distributed Stochastic Neighbor Embedding (t-SNE) was applied using correlation distance metrics.

